# Immunogenicity and Efficacy of TNX-1800, A Live Virus Recombinant Poxvirus Vaccine Candidate, Against SARS-CoV-2 Challenge in Nonhuman Primates

**DOI:** 10.1101/2023.09.19.558485

**Authors:** Mayanka Awasthi, Anthony Macaluso, Dawn Myscofski, Jon Prigge, Fusataka Koide, Ryan S Noyce, Siobhan Fogarty, Helen Stillwell, Scott J Goebel, Bruce Daugherty, Farooq Nasar, Sina Bavari, Seth Lederman

**Author notes:** Correspondence: Seth Lederman.

## Abstract

TNX-1800 is a synthetically derived live chimeric Horsepox Virus (rcHPXV) vaccine expressing Wuhan SARS-CoV-2 spike (S) protein. The primary objective of this study was to evaluate the immunogenicity and efficacy of TNX-1800 in two nonhuman primate species challenged with USA-WA1/2020 SARS-CoV-2. TNX-1800 vaccination was well tolerated, as indicated by the lack of serious adverse events or significant changes in clinical parameters. A single dose of TNX-1800 generated robust humoral responses in African Green Monkeys and Cynomolgus Macaques, as measured by the total binding anti-SARS-CoV-2 S IgG and neutralizing antibody titers against the USA-WA1/2020 strain. In Cynomolgus Macaques, a single dose of TNX-1800 induced a strong interferon-gamma (IFN-γ) mediated T cell response, promoting both pathogen clearance in the upper and lower airways and generation of systemic neutralizing antibody response against WA strain SARS-CoV-2. Future studies will assess the efficacy of TNX-1800 against newly emerging variants and demonstrate its safety in humans.

## Introduction

As the SARS-CoV-2 infection continues to evolve [1], the urgency to develop and distribute more effective, long-lasting vaccines against continually emerging SARS-CoV-2 variants remains a top priority for global public health [2]. Despite significant progress, challenges persist in ensuring equitable vaccine access and distribution [3]. Vaccines have proven to be cost-effective compared to other medical interventions, benefiting from established technology, mass production, simplicity, government support, and global health initiatives [4]. Their potential to prevent disease spread and reduce long-term healthcare burdens further emphasizes their significance [5-7].

Currently, there are several SARS-CoV-2 vaccines that are authorized by the U.S. Food and Drug Administration (FDA) [8, 9]. Pfizer-BioNTech and Moderna COVID-19 vaccines which are mRNA vaccines [10] and Novavax COVID-19 vaccine which is a protein subunit vaccine [11]. J&J/Janssen COVID-19 vaccine [12], a viral vector vaccine has expired and is no longer available for use in the United States as of May 6, 2023 [13]. The most used vaccines share a common technology, utilizing mRNA to deliver genetic instructions to produce the SARS-CoV-2 spike protein within the body [14]. All the available vaccines are based on multiple dosing, and some require specific storage and transportation at very cold temperatures [15].

Compared to many other vectors, poxvirus vectors offer several advantages, including potential logistical benefits [16]. Poxvirus does not require ultra-cold storage conditions, simplifying distribution in resource-limited settings [17-19]. Our recent study showed that a single dose vaccination with TNX-801, a recombinant chimeric horsepox virus (rcHPXV), was effective at protecting NHPs from infection with monkeypox virus (MPXV) [20].

TNX-1800, a synthetic-derived live chimeric Horsepox Virus (rcHPXV) vaccine comprising an engineered SARS-CoV-2 S (spike) gene expression cassette (Figure 1). TNX-1800 incorporates the spike glycoprotein gene sequence from the SARS CoV-2 virus (Wuhan strain, GenBank NC_045512) into the HPXV genome, and have the potential to overcome pre-existing immunity, address concerns related to adverse events associated with certain vector platforms, and induce a stronger immune response making them promising candidates in advancing global vaccination efforts against COVID-19 and other infectious diseases [21].

**Figure 1.**
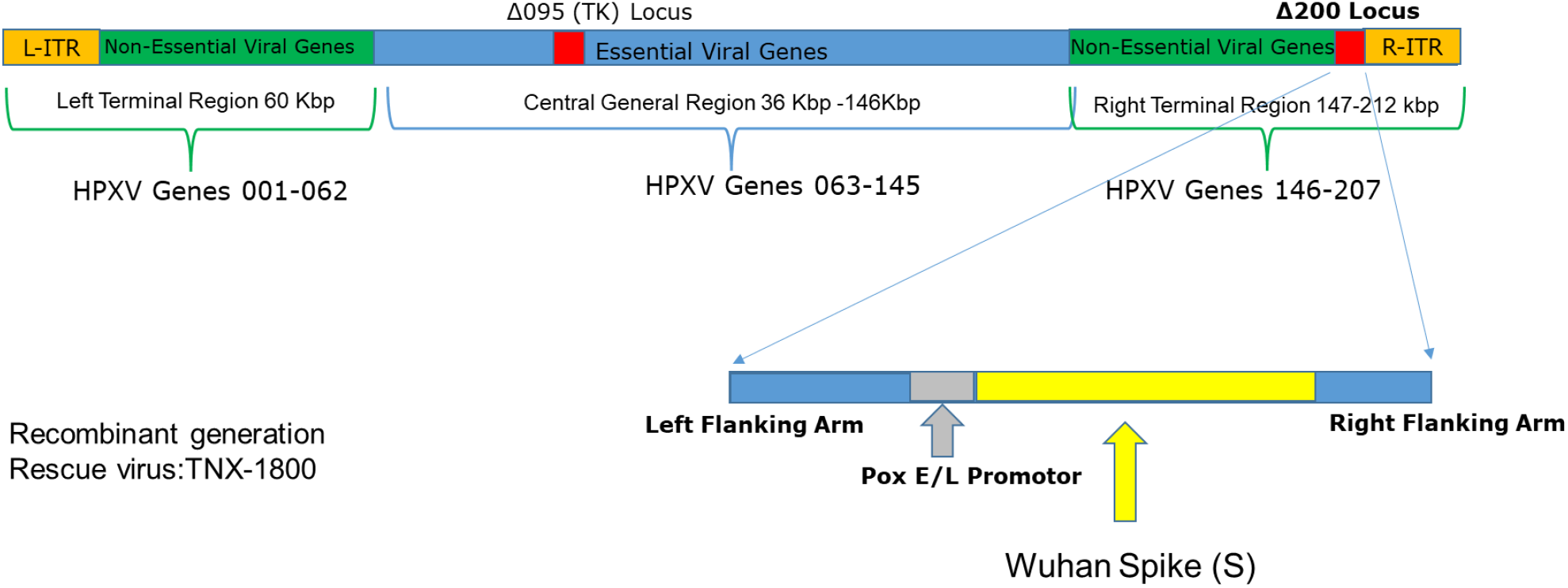
Schematic of the recombinant TNX-1800 generation. TNX-1800 was constructed by homologous recombination, inserting the SARS-CoV-2 spike gene (codon optimized for VACV and under the control of a synthetic promotor) into the Δ200 locus of the recombinant chimeric Horsepox Virus (rcHPXV). The resulting construct contains Pox E/L promoter-driven expression fragments for Spike Glycoprotein armed with left and right HPXV flanking regions.

The recombinant modified vaccinia virus Ankara (MVA) vaccine expressing a prefusion-stabilized SARS-CoV-2 spike glycoprotein induces robust immunity and protection in mouse and hamster model [22]. A modified MVA has also been shown to protects against SARS-CoV-2 in Syrian hamsters and non-human primates [23]. We previously have assessed TNX-1800 immunogenicity, tolerability, and vaccine systemic dissemination in Syrian golden hamsters and New Zealand white rabbits [21]. In this proof-of-concept report, we have investigated the immunogenicity and efficacy of TNX-1800 in nonhuman primates. The work presented in this report demonstrates vaccine efficacy by a rcHPXV construct expressing the Wuhan strain SARS-CoV-2 spike proteins. The ability of TNX-1800 to confer broad cross-protective immunity against newly emerging variants will require further investigation.

Here, TNX-1800 has been evaluated in African Green Monkeys (AGMs) and Cynomolgus Macaques (CMs), challenged with the SARS-CoV-2 virus (WA1). In both NHP species, TNX-1800 vaccination was well tolerated, indicated by the lack of serious adverse events or significant changes in clinical parameters. A single dose of TNX-1800 was able to generate robust humoral responses in AGMs, as measured by total binding IgG and neutralizing antibody titers. In CMs, a dose of 1.3 x 10^6^ PFU of TNX-1800 elicited a strong T-cell response, promoting both pathogen clearance and the generation of neutralizing antibody responses against WA1 SARS-CoV-2.

## Materials and Methods

### Vaccine design

TNX-1800 was constructed by homologous recombination, inserting the SARS-CoV-2 spike gene (codon optimized for vaccinia virus and under the control of a synthetic promotor) into TNX-801 at the Δ200 gene locus. Diluent was a sterile solution of Tris-HCl 10 mM; pH 8.0).

### Ethics Statement

This work was supported by an approved Institute Animal Care and Use Committee (IACUC) animal research protocol. The research was conducted under an IACUC-approved protocol in compliance with the Animal Welfare Act, PHS Policy, and other Federal statutes and regulations relating to animals and experiments involving animals. The facility where this research was conducted is accredited by the Association for Assessment and Accreditation of Laboratory Animal Care (AAALAC International) and adheres to principles stated in the Guide for the Care and Use of Laboratory Animals, National Research Council, 2011.

### Study Design and Immunization Procedure

A total of 8 African Green Monkeys (AGM) and 8 Cynomolgus Macaques of Mauritius origin (CM) were purchased from vendors Primera Science Center (LaBelle, FL 33935) and PrimGen (Hines, IL), respectively. Animals were subsequently quarantined in the ABSL-2 at the Piccard facility for a minimum of 42 days. Prior to release from quarantine, the animals were free of obvious abnormalities indicative of any health problems and were confirmed to be free of simian immunodeficiency virus (SIV), Simian Retrovirus (SRV)-1/2/3/4/5, simian T-lymphotropic viruses (STLV)-1, Measles, and Tuberculosis (TB). Considering the day of vaccination as day 0 of the study, all animals were challenged with SARS-CoV-2-WA (Washington strain) on day 41 of the study via the IN/IT route. Samples for different assessments were collected on the days indicated in **Figure 2A** and **Figure 3A**.

**Figure 2.**
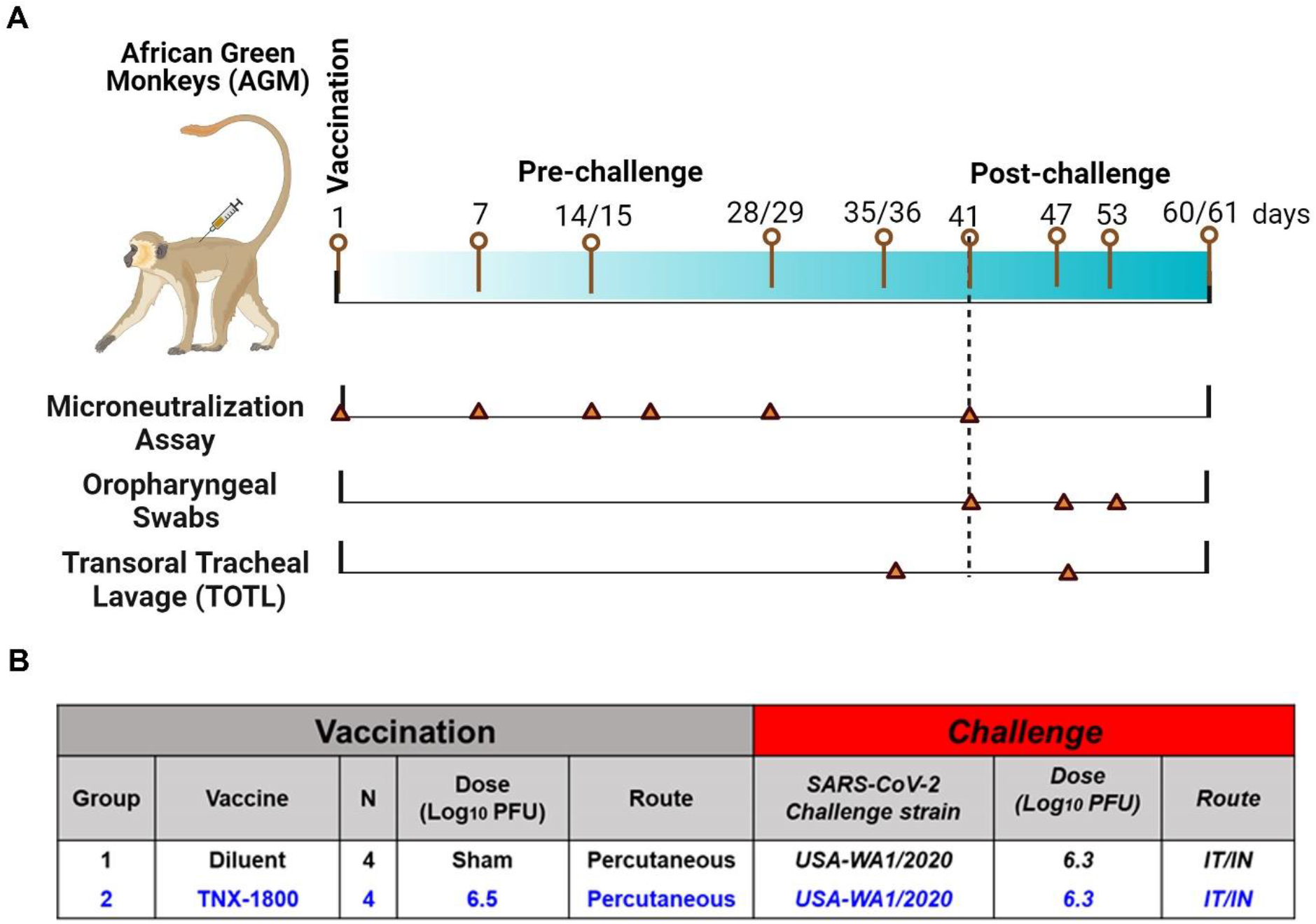
Study design for immunogenicity and efficacy of TNX-1800 in African Green Monkeys. (A) Study schedule for collecting serum samples from monkeys vaccinated with TNX-1800 and challenged with the SARS-CoV-2 isolate, USA-WA1/2020 on day 41 post vaccination. (B) TNX-1800 dosing concentrations and dose administration routes adopted for pre- and post-challenge immunogenicity and efficacy study.

**Figure 3.**
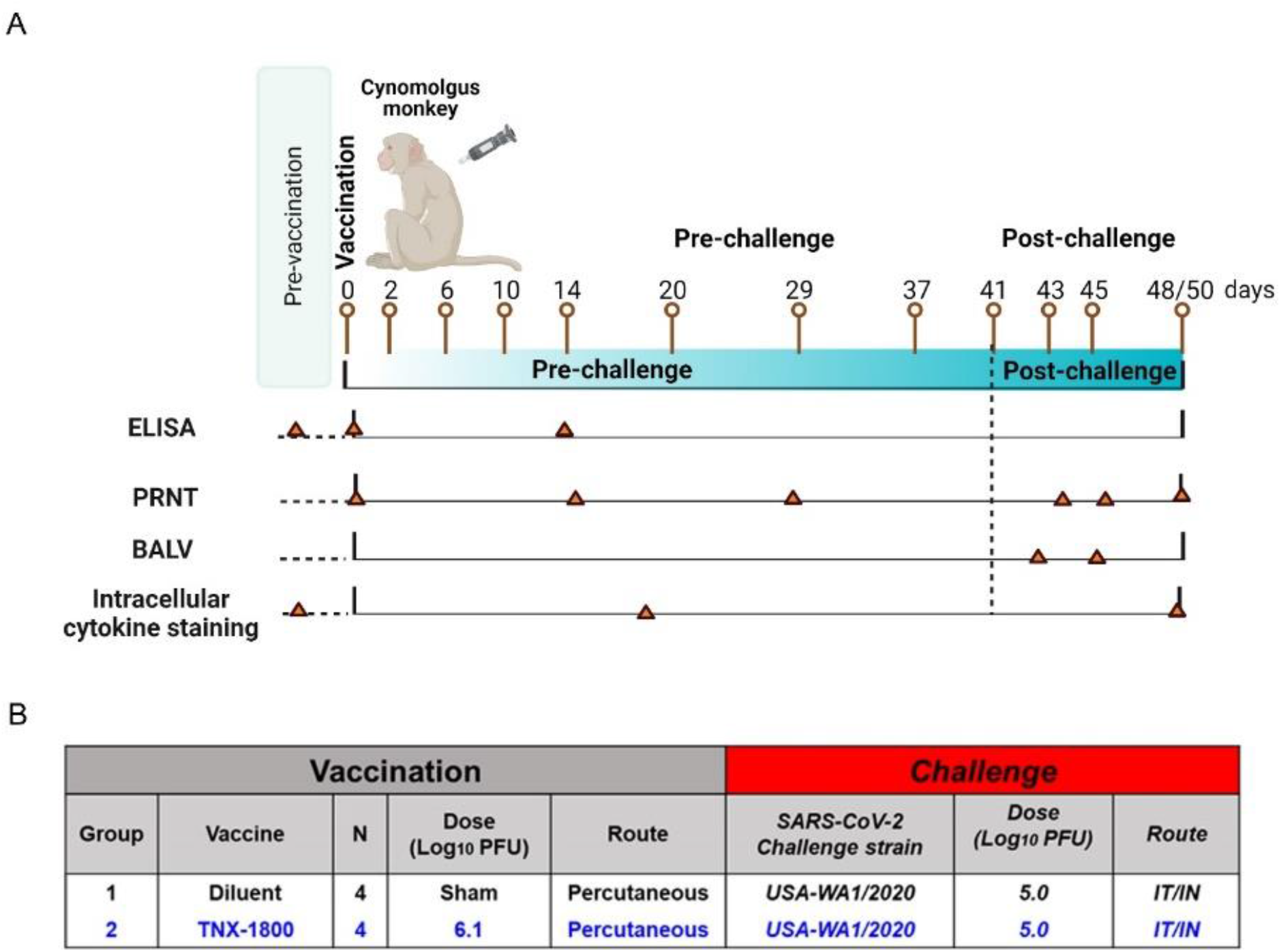
Study design for immunogenicity and efficacy of TNX-1800 in Cynomolgus monkeys. (A) Schedule for pre- and post-vaccination serum collection from monkeys vaccinated with TNX-1800 and challenged with the SARS-CoV-2 isolate, USA-WA1/2020 on day 41 post vaccination. (B) TNX-1800 dosing concentrations and dose administration routes adopted for pre- and post-challenge immunogenicity and efficacy study in CM.

For the AGM study, four naïve monkeys (2 male and 2 female) were vaccinated with 2.9 x 10^6^ PFU of TNX-1800 (10 μL total volume) via scarification at the interscapular region on Day 0. Similarly, for the CM study, four adult monkeys (2 male and 2 female) were vaccinated with 1.3 x 10^6^ PFU of TNX-1800 (2.5 μL total volume) via scarification at the interscapular region on Day 0. For the control groups of each study, four naïve monkeys (2 male/2 female) were injected with a sterile solution of Tris-HCL (10 mM; pH 8.0). Following vaccine administration, a bifurcated needle was used to scarify the area by penetrating the skin vertically approximately 15 times.

For both studies, clinical signs, body weight and body temperature were assessed throughout the study at predetermined time points. For the AGM study, oropharyngeal (OP) swabs and Transoral Tracheal Lavage (TOTL) were collected according to the schedule indicated in **Figure 2A**: Days 35/36 (TOTL only), day 41 (OP only), days 47/48, 53 and 60/61 for RT-qPCR (quantifying SARS-CoV-2 viral genomes). PBMCs were collected for cytokine quantification. Animals were euthanized on day 47 (6 days post-infection; 1/sex/group) or 60 (19 days post-infection; 1/sex/group). Histopathology was completed on cranial and caudal lung lobes.

For the CM study, blood collection occurred on days indicated in **Figure 3B** and were used for the measurement of binding and neutralizing antibodies through enzyme-linked immunosorbent assay (ELISA) and plaque reduction neutralization (PRNT) assay. PBMCs were also isolated for flow cytometry analysis and cytokine quantification. Nasal swabs and bronchoalveolar lavage (BALV) samples were collected on days 42 through 45 and days 42/45, respectively. All animals were euthanized either on day 48 (7 days post-infection; 1/sex/group) or day 50 (9 days post-infection; 1/sex/group).

### SARS-CoV-2 Challenge and Back-Titration of Inoculum

The animals were anesthetized intramuscularly with 10 mg/kg of Ketamine. Virus inoculum was prepared from the stocks and diluted in sterile PBS to a dose level of 1x10^5^ TCID_50_ per 2 mL volume. Syringes of 2 mL volume were prepared for each animal. The animals were challenged via the intranasal route and followed by the intratracheal route. For the intranasal route, 0.5 mL of the viral inoculum was administered dropwise into each nostril for a total of 1 mL volume per animal. Upon administration, the animal’s head was tilted back for about 20 seconds. The animal was returned to its housing unit and monitored until fully recovered. For the intratracheal route, 1 mL of diluted virus was delivered intratracheally using a French rubber catheter/feeding tube, size 10, sterile, (cut 4”-6” in length). The syringe containing the inoculum was attached to a sterile French catheter or feeding tube. The small end of the feeding tube with inoculum was inserted into the glottis. Once in place, the inoculum was injected into the trachea and then the catheter was removed from the trachea. New or sterilized equipment was used for each animal. The study animal was then inoculated via the intranasal route. The remaining inoculum was aliquoted into cryovial tubes and stored at -70oC or below. The stored challenge inoculum was later back tittered via a TCID_50_ assay for verification of proper dose level.

### Enzyme-linked Immunosorbent Assay (ELISA)

A standard indirect ELISA was performed to analyze serum samples for binding antibodies to the SARS-CoV-2 spike protein. For this assay, 96-well plates were coated with 50 μL of SARS-CoV-2 spike protein (Sino Biological, Beijing, China) diluted to 2 μg/mL in Carbonate-Bicarbonate Buffer (CBB, Sigma-Aldrich, St. Louis, MO). Plates were incubated overnight at 2-8 oC. Unbound coating antigen in each well was removed by washing with PBS + 0.05% Tween-20 and then blocked with PBS + 1% BSA. Test and positive control samples were diluted in assay diluent (PBS-Tween 20; 1% BSA) to a starting point dilution of 1:20, followed by four-fold serial dilutions. Once blocking was completed, the Blocking Buffer was removed by inversion and each sample was plated. Plates were incubated for 60 to 70 minutes at room temperature, followed by washing with PBS + 0.05% Tween-20 to remove unbound sera. Secondary detection antibody (Goat anti-Monkey IgG (H+L) Secondary Antibody, HRP, Invitrogen, Carlsbad, CA) was added at a dilution of 1:10,000, and plates were incubated for 60 to 70 minutes at room temperature. Unbound antibodies were subsequently removed by washing with PBS + 0.05% Tween-20 and a final wash with PBS alone. To develop, 1-Step Ultra TMB substrate (SeraCare Life Sciences, Gaithersburg, MD) was added to each well and the plate was developed. The reaction was stopped after 10 minutes to 15 minutes with TMB stop solution (SeraCare Life Sciences). The plates were read within 30 minutes at 450 nm using a Thermo Lab Systems Multiskan spectrophotometer.

### Plaque Reduction Neutralization (PRNT) Assay

Serum was isolated and aliquoted into a minimum of two single-labeled cryovial tubes of 200 μL each. A VACV-specific PRNT assay was used to determine if the vaccinations have elicited a neutralizing antibody titer against HPXV. Serum samples were heat inactivated for 30 minutes at 56°C. Briefly, serum samples were serially diluted in media and added to equal volume of a fixed dilution of the Western Reserve strain of VACV. The serum-virus mixture will then be incubated for 1 hour at 37°C. Subsequently, 250 μL of each serum-virus mixture will be added in duplicate to a near confluent monolayer of Vero cells and incubated at 37 ± 2°C; 5% CO_2_ for 1 hour. Afterwards, 1 mL of 0.5% methylcellulose in DMEM + 2% FBS was added to each well and incubated at 37 ± 2°C, 5% CO_2_ for 3 days. The virus-cell mixture was incubated at 37 ± 2°C; 5% CO_2_ for 48 ± 4 hours, fixed with cold methanol and then stained with 0.2% Crystal Violet solution for the enumeration of the plaques. Neutralization end-point titers were calculated and based on the reciprocal dilution of the test serum that produced 50% plaque reduction compared to the virus only control (PRNT_50_).

### Intracellular Cytokine Staining for SARS-CoV-2-Specific Immune Responses

Intracellular cytokine staining via flow cytometry was performed from cryopreserved PBMCs. Cells were thawed in media containing >5 U/mL of benzonase and resuspended in complete RPMI media supplemented with 10% FCS, L-Glutamine and Penicillin– Streptomycin at a concentration of 2x10^7^ cells/mL. Then, 1x10^6^ PBMCs per well, was plated in a 96-well plate and stimulated with synthetic peptides spanning the SARS-CoV-2 spike protein. One well per NHP was stimulated with 10 pg/mL phorbol myristate acetate (PMA) and 1 μg/mL ionomycin (Sigma-Aldrich) (P+I) as a positive control. PBMCs was co-stimulated in the presence of anti-human CD28, CD49d (1 μg/ ml; Life Technologies) and CD107a (BioLegend) then incubated for 16 hours (± 2 hours) after the addition of 1 μg/ ml of brefeldin A and Monensin to each well. The antibody panel included live/dead dye for 10 mins and then stained with predetermined titers of mAbs against CD4, CD8, CD3 for 30 min intracellularly with mAbs against IFN-γ and IL-4. Samples were acquired on BIOQUAL MACSQuant Analyzer 16. Analysis was performed using FlowJo Software (Version 10).

### Collection of Oropharyngeal Swabs

The animals were anesthetized prior to the swab procedure. For oropharyngeal swab the animal’s mouth was cleaned of excess saliva and food particles. The swab was inserted into the animal’s mouth taking care not to touch any surface of the mouth until the tip of the swab was past the base of the tongue. Once there, the swab was pressed against the dorsal surface of the pharyngeal area and rolled across the surface. The swab was removed, taking care to avoid touching any other mouth surfaces. Following collection, the swabs were placed into a collection vial (2/specimen).

### Collection of Transoral Tracheal and Bronchioalveolar Lavage

The Transoral Tracheal Lavage (TOTL) procedure was performed on the anesthetized animal by inserting a tube into the trachea. Once the end of the tube was situated approximately at the mid-point of the trachea, a syringe containing up to 5 mL of sterile PBS was attached to the tube, and the medium was slowly instilled into the trachea. Once the instillation was complete, negative pressure was immediately applied via the same syringe to collect as much of the PBS as possible. Samples were snap-frozen immediately following collection and stored at or below -70 °C until analyzed.

The bronchoalveolar lavage (BAL) procedure was performed on the anesthetized animal by the “chair method”. The NHP was placed in dorsal recumbency in the chair channel such that the handle was tightened into the position supporting the animal’s weight. A catheter tube was premeasured and marked before placement. The animal’s head was tilted back and down below the edge of the chair channel. The catheter tube was then inserted into the animal’s trachea via a laryngoscope during inspiration. The tube was placed into the correct positioning and the BAL wash procedure was executed. A total of 10 mL was flushed through the tube. Upon sample completion, the handle at the base of the chair was loosened and the animal was removed from the chair. If any complications occur, the procedure was stopped, and the clinical veterinarian was consulted for treatment. The volume instilled and recovered from each animal, as well as any presence of blood in the BAL samples, was recorded. The animal was monitored until fully recovered and for the first hour post-BAL collection. If warranted, additional monitoring was also conducted to ensure animal health. The collected BAL samples were placed immediately onto wet ice and processed for isolation of fluid. The sample was centrifuged to pellet debris, supernatant collected, and then aliquoted.

### Viral Load RT-qPCR

Viral loads in swab samples were measured using RT-qPCR. Total DNA was extracted from each sample using QIAcube robot according to manufacturer protocol and viral genome copies were quantified via qPCR. Briefly, samples were extracted using TriZol reagent (Sigma Aldrich) following manufacturer’s recommendations. Viral DNA isolated from samples was eluted with nuclease free H_2_O and stored at -70°C or below. The quantitative PCR (qPCR) reaction used to assess vaccine viral load, was targeted to the Orthopox E9L DNA Polymerase gene using commercially available Pan-Orthopox Virus E9L Gene-Specific Quantitative PCR Assay Detection Kit (BEI Resources Cat# NR-9350).

### Infectious Viral Load Assay (TCID_50_)

Vero TMPRSS2 cells (obtained from the Vaccine Research Center-NIAID, Rockville, MD) were plated at 25,000 cells/well in DMEM + 10% FBS + Gentamicin and the cultures are incubated at 37°C, 5.0% CO2. Cells were 80 -100% confluent the following day. Medium was aspirated and replaced with 180 μL of DMEM + 2% FBS + gentamicin. Twenty (20) μL of the sample was added to the top row in quadruplicate and mixed with a pipettor. Using the pipettor, 20 μL was transferred to the next row and repeated down the plate (columns A-H) representing 10-fold dilutions. The tips were disposed of for each row and repeated until the last row. Positive (virus stock of known infectious titer in the assay) and negative (medium only) control wells were included in each assay set-up. The plates were incubated at 37°C, 5.0% CO_2_ for 4 days. The cell monolayers were visually inspected for cytopathic effects (CPE). Non-infected wells had a clear confluent cell monolayer while infected wells had little cell presence. The presence of CPE was marked on the lab form as a “+” and absence of CPE as “0”. The TCID_50_ value was calculated using the Read-Muench formula. For optimal assay performance, the TCID_50_ value of the positive control should test within 10-fold of the expected value.

### Statistical Analysis

All statistical analyses were performed in GraphPad Prism (v9) by ordinary one-way Anova followed by multiple comparisons between groups*; p < 0.05 against non-treated group ****; p < 0.0001 against the non-treated group.

## Results

### African Green Monkeys (AGMs) Challenge study

#### Study TNX-1800 elicited robust immune responses against SARS-CoV-2 in AGMs

TNX-1800 immunization was well-tolerated in all monkeys and no serious adverse events were observed. All four animals vaccinated with TNX-1800 showed measurable vaccine “take”, or cutaneous reaction indicating successful vaccination and all animals seroconverted as early as day 14 (suggests 100% seroconversion). As shown in **Figure 4A**, the magnitude of the neutralizing antibody response elicited was sufficiently high, and group geometric mean titers (GMT) detected on the day of challenge (Day 41) were 80. On Day 41, animals were challenged with US COVID-19 isolate WA1 strain through intranasal and intratracheal routes. Viral shedding in the upper respiratory tract (URT) was assessed by OP swabs and viral load in the lower respiratory tract (LRT) by TOTL, using qRT-PCR to determine the number of genome copies of SARS-CoV-2 present in the samples (Figure 4B and C). All the animals vaccinated with TNX-1800 showed no measurable viral RNA in URT, six days post-challenge. As presented in **Figure 4B**, a significant reduction of viral genome copies was observed in transoral-tracheal (TOTL) lavage by TNX-1800 compared to the diluent control (*p=0.0328). The TNX-1800 treated animals didn’t show significant viral load (>1000 copies/mL) in TOTL whereas animals in control groups exhibited peak individual values ranging from 1.17E+03 to 2.11E+06 copies/mL, when observed six days post-challenge. As presented in **Figure 4C**, a strong reduction in SARS-CoV-2 genome copy number in OP mucosa was observed in the group vaccinated by TNX-1800 compared to the control groups (*p=0.0285). These data suggest that protection was successfully induced in both the LRT and URT.

**Figure 4.**
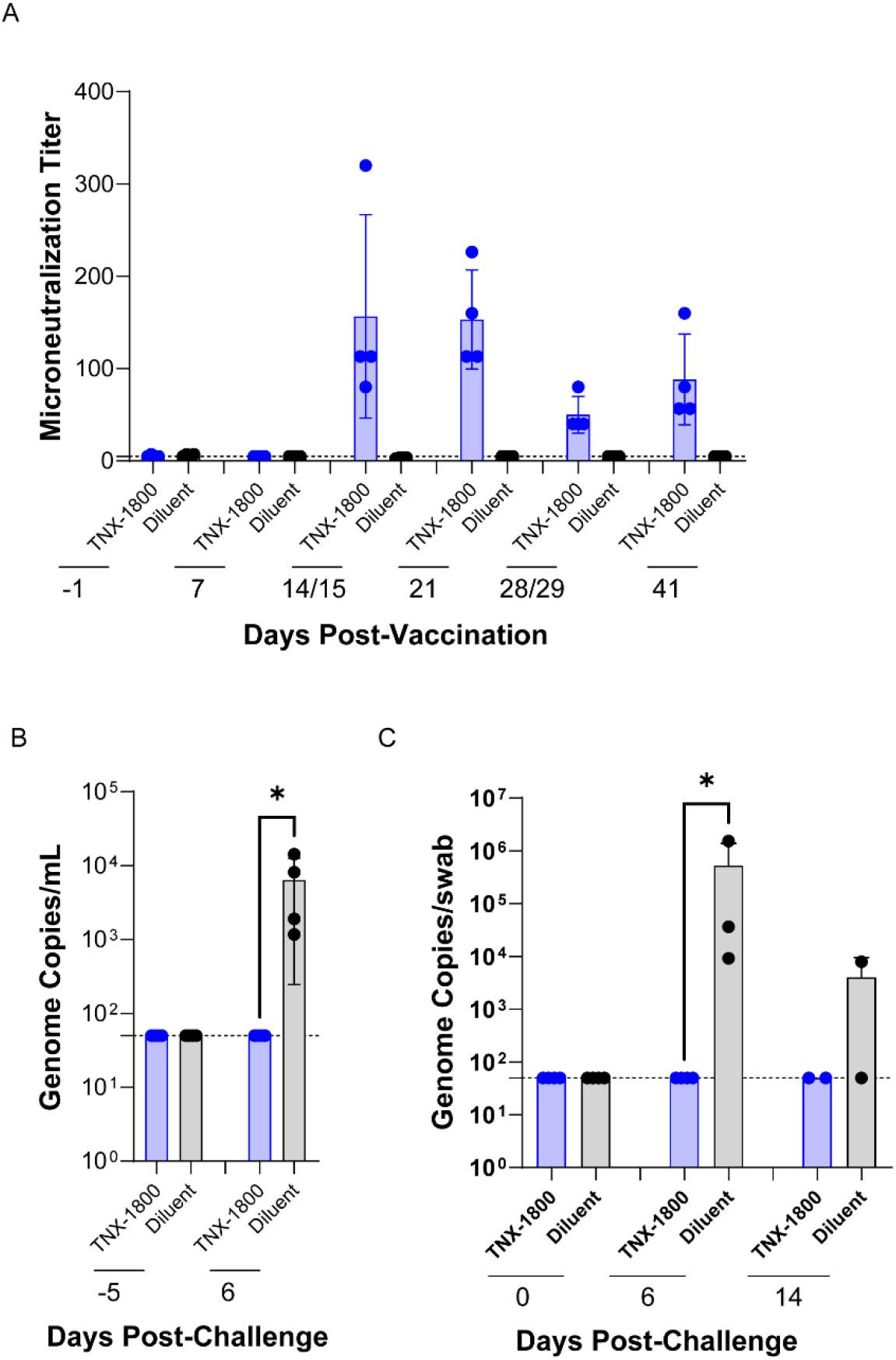
TNX-1800 immunogenicity and post-challenge viral shedding in African Green Monkeys. (A) Neutralizing antibody titers against SARS-CoV-2 were determined via microneutralization assay in serum samples collected at selected time points of the study. (B) SARS-CoV-2 load quantified in AGM tracheal lavage samples by RT-qPCR. Significant reduction of SARS -CoV2 shedding was observed in transoral-tracheal (TOTL) lavage by TNX-1800 (*p=0.0328). (C) SARS-CoV-2 load quantified in AGM oropharyngeal (OP) swabs by RT-qPCR. Strong reduction in SARS-CoV-2 shedding in oropharyngeal (OP) mucosa was observed in the group vaccinated by TNX-1800 compared to the control groups (*p=0.0285).

There were no treatment-related differences in macroscopic or microscopic observations. The chronic inflammation was minimal to mild and appeared similarly across both groups. The adhesion and inflammation was observed in the left caudal lung of one animal of the control group and was most likely related to an amorphous foreign body than challenge with SARS-COV-2.

### Cynomolgus Monkeys (CMs) Challenge Study

#### Humoral Immune Responses in CMs

Animals were observed and scored as alert and healthy for the duration of the study, and no adverse events or significant changes in clinical parameters were recorded. The serum titers of SARS-CoV-2 spike antigen reactive IgG antibodies in all animals were measured at days 0 and 14 after vaccination and compared with pre-vaccination values (**Figure 5A**). In the TNX-1800 dose group, a mean endpoint binding titer of 5319 against the SARSCoV-2 spike S1+S2 specific isotype IgG1 titer was measured 14 days after vaccination. The pre-vaccination IgG1 titer value for this group was 95. These results indicated that the level of SARS-CoV-2 neutralizing antibodies in the serum of the animals vaccinated with TNX-1800 was significantly higher as compared to the diluent control group. In summary, the percutaneous delivery of TNX-1800 induced a functional T-cell response and SARS-CoV-2 spike protein-directed humoral immunity.

**Figure 5.**
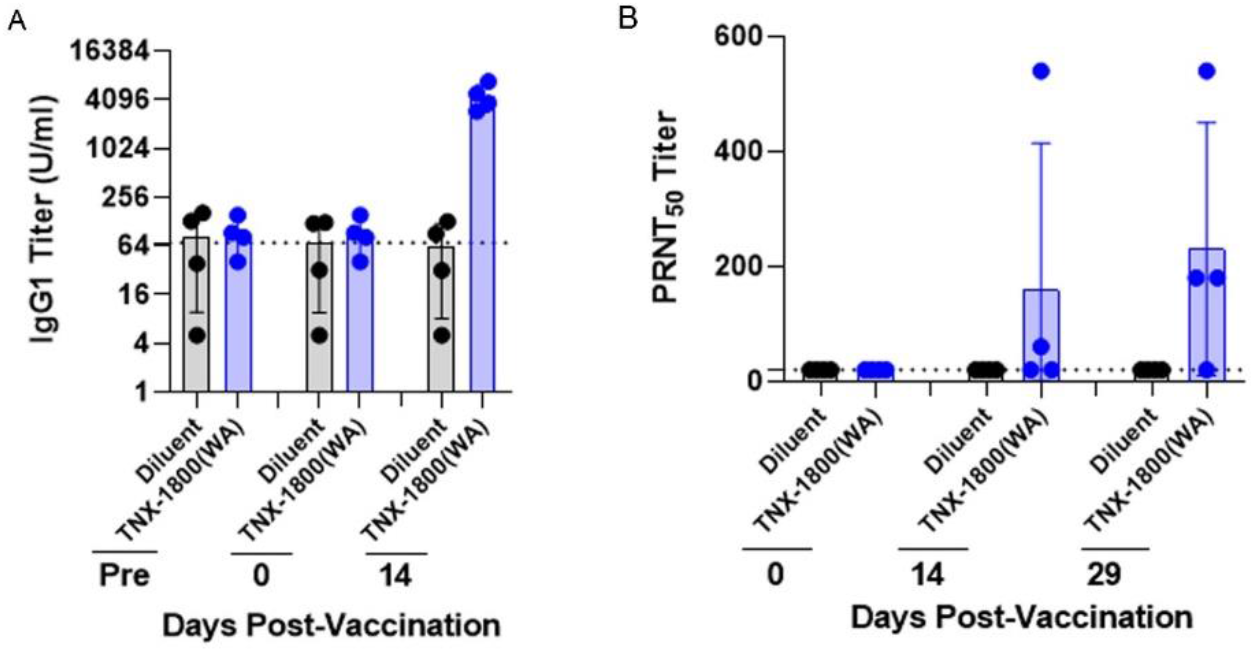
Total and neutralizing antibody titers against SARS-CoV-2 in TNX-1800 vaccinated Cynomolgus Monkeys. (A) Sera collected were analyzed in an ELISA for binding total IgG antibody levels to the SARS-CoV-2 Spike antigen comprised of three peptide fragments. The range of dilutions for the sera ranged from a reciprocal dilution of 50 to 6400 consisting of 4-fold serial dilutions. The stippled lines indicate the dilution of 1:64. Each data point depicts the individual animal, and the bar represents the group mean of the difference between OD450 and OD630. (B) The sera from select timepoints outlined in Figure 2 were processed and analyzed for neutralizing antibody titers against the challenge virus, USA-WA1/2020. The range of dilutions for the sera ranged from a reciprocal dilution from 20 to 4860 consisting of 3-fold dilutions. The stippled line indicates the minimum dilution of 1:10. Each data point depicts the individual animal.

To evaluate the humoral immune response elicited by the TNX-1800 vaccination both pre and post SARS-CoV-2 challenge, the PRNT assay was performed (**Figure 5B**). Within 14 days post vaccination, 50% of the TNX-1800 vaccinated animals exhibited elevated neutralizing antibody responses, from respective baseline values with an increase in PRNT_50_ titers ranging from 3- to 27-folds. Further, all the animals exhibited 3- to 243-folds elevation in PRNT_50_ titers comparable to respective baseline values 29 days post vaccination.

#### Cell-Mediated Immune Responses

Intracellular cytokine staining (ICS) by flow cytometry was performed in all the CM. animals at days -174, -12, and 7 post-vaccination (**Figure 6**). In most of the animals, both CD4+ and CD8+ effector T-cells showed elevated levels of interferon-gamma (IFN-γ) on day 7 post-challenge in comparison to the levels prior to the challenge (**Figure 6A and 6C**). However, both CD4+ and CD8+ T-cells showed no significant levels of changes in interleukin 4 (IL-4) levels at any time point (**Figure 6B and 6D**). This shows that the immune response in the TNX-1800 vaccinated animals post SARS-CoV-2 exposure was most likely an IFN-γ mediated cytokine response.

**Figure 6.**
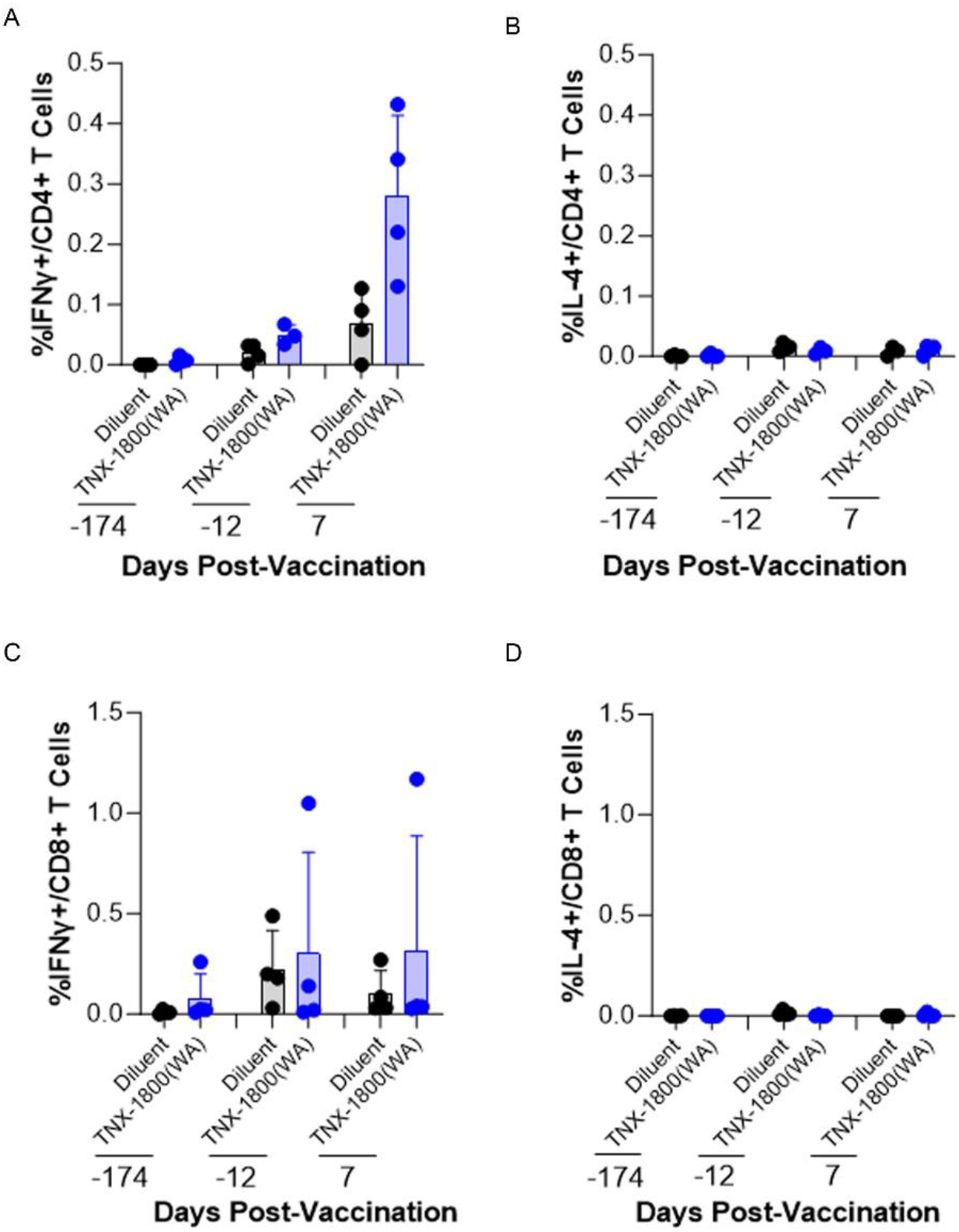
SARS-CoV2 specific cell mediated immune response to TNX-1800 vaccination in Cynomolgus Monkeys. (A-D) Relative distribution of Intracellular Cytokines in PMBCs. PBMCs were collected and cryopreserved and then intracellularly stimulated with spike antigens and stained with monoclonal antibodies against IL-4 and IFN-γ at selected time points. The scale of the vertical axis is specific to each analyte and corresponds to the median percentage of positive cells.

Viral loads in the upper and lower respiratory tracts after SARS-CoV-2 challenge were assessed by Tissue Culture Infectious Dose (TCID) assay. The TCID assay for BALF showed the detection of post challenge viral strains in most animals, with infectious titers quantified between about 104 and 107 TCID_50_/ml. The TCID data of BAL samples revealed the significant reduction in the TCID_50_ values 2 days post-inoculation (dpi) in TNX-1800 vaccinated animals after SARS-CoV2-WA-1 challenge. Analysis of viral loads in, at 2 dpi showed the reduced viral load levels in TNX-1800 with respect to diluent control (**Figure 7**), which further ensures the efficacy of TNX-1800 over SARS-CoV-2-WA-1 challenge in all the animals.

**Figure 7.**
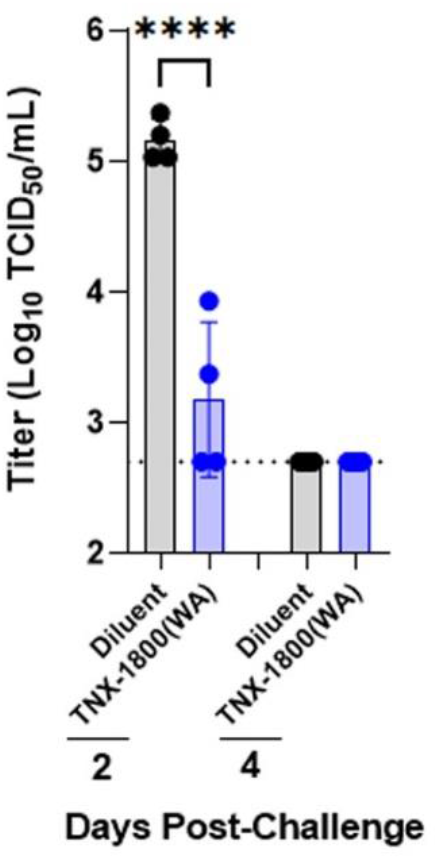
Infectious Viral Loads in BAL samples from SARS-CoV-2 Challenged Macaques. BAL samples from SARS-CoV-2 challenged monkeys were collected and tested for amounts of infectious virus loads by TCID50 assay. Each data point depicts the individual animal. Significant reduction (*p<0.0001) in infectious viral load was observed in the vaccinated group. The stippled line represents the lower limit of detection (2.70 Log TCID_50_/mL).

## Discussion

As the world continues to combat the SARS-CoV-2 infection, ongoing research and innovation are essential to develop additional safe and accessible vaccines [24]. While currently approved vaccines have played a significant role in saving lives, newer platforms particularly ones that are easier or cheaper to manufacture and deliver are needed to get control of this pandemic. The poxvirus vectors show great potential to address existing limitations and contribute to achieving global vaccination goals. Previously, in two animal models, the Syrian hamsters and New Zealand white rabbits [21], we demonstrated tolerability, safety, and immunogenicity of TNX-1800. In the current report, based on two NHP models, we confirmed the immunogenicity and efficacy of TNX-1800, a live virus recombinant poxvirus vaccine candidate against the SARS-CoV-2 challenge.

The work presented in this report demonstrates vaccine efficacy of TNX-1800 in NHP species. TNX-1800 vaccination using percutaneous method was well-tolerated, with no reports of serious adverse events or significant changes in clinical parameters in two NHP models. Thus, providing the possible use of alternative routes to deliver the vaccine effectively. Remarkably, a single dose of TNX-1800 administered to NHPs resulted in robust humoral responses, as indicated by elevated levels of total SARS-CoV-2 S binding total IgG and neutralizing antibodies compared to the diluent control. The neutralizing antibody responses induced by TNX-1800 were associated with upper and lower respiratory tract protection against mucosal challenge, meeting the primary immunogenicity and efficacy endpoints.

For CMs, a specific dose of TNX-1800 (1.3 x 10^6^ PFU) elicited a strong T-cell response, promoting both pathogen clearance and the generation of long-lasting neutralizing antibody responses against SARS-CoV-2. The observed elevation of interferon-gamma (IFN-γ) levels in CMs vaccinated with TNX-1800 indicates a predominant TH1 immune response. IFN-γ is a pro-inflammatory cytokine crucial for antiviral defense, inhibiting viral replication and enhancing the activity of cytotoxic T cells and natural killer (NK) cells. This suggests that TNX-1800 effectively primed T cells to recognize SARS-CoV-2-infected cells and mount a robust cellular immune response to clear the virus from the body.

Noteworthy is the absence of significant changes in interleukin-4 (IL-4) levels in CMs post-SARS-CoV-2 challenge, indicating that the immune response primarily involved a TH1 immune response driven by IFN-γ rather than a TH2 response associated with specific antibody production by B cells. This balanced immune response is essential in combating decreasing unwanted toxicity, viral infections, as cellular immunity, including T-cell responses, plays a crucial role in eliminating virus-infected cells.

In conclusion, the cytokine profiling assay demonstrated that TNX-1800 vaccination in CMs induced a strong TH1 response, promoting pathogen clearance and generating long-lasting neutralizing antibody responses against SARS-CoV-2. The vaccine’s ability to elicit a comprehensive immune response involving both antibody-mediated and cellular (T cell) immunity contributes to its effectiveness in promoting pathogen clearance and generating long-lasting protection against SARS-CoV-2 infection. Additionally, the significant reduction of viral shedding in the upper airway and bronchoalveolar lavage indicates that TNX-1800 induced a protective mucosal immune response, which is critical in preventing infections at the entry sites of respiratory viruses and may also play a role in decreasing transmission.

In NHPs, the HPXV backbone platform (TNX-801) was shown to be well tolerated, immunogenic, and provided complete protection against the lethal monkeypox challenge [20]. The engineered monovalent vector expressing SARS-CoV-2 spike protein demonstrated a similar safety and efficacy profile. Taken together, these data strongly suggest that the HPXV can be developed into a modular “plug and play” multi-valent platform to provide efficacy against multiple pathogens.

Most licensed vaccines are administered by intramuscular or subcutaneous routes. These mechanisms of vaccination may not appropriately prime the mucosal lymph nodes or produce sufficient IgA responses to provide durable protection against SAR-CoV-2 mediated disease. In contrast, intradermal vaccination has the potential to generate respiratory, secretory IgA (sIgA) and memory mucosal T-cell responses. In fact, vaccination via intradermal routes has been shown to induce skin Langerhans cells to present their antigenic cargo in mucosal draining lymph nodes to mucosal T cells which allows mucosal T-cell memory generation and robust long-lasting sIgA responses [25]. Although we have not formally tested neutralizing sIgA in the respiratory system after vaccination, the NHP data from oropharyngeal swabs, TL, and BAL firmly suggests the presence of a strong humoral response at mucosal sites after a single vaccination. Currently, studies are being conducted to investigate the presence of spike-specific neutralizing antibodies at the mucosal sites following TNX-1800 vaccination.

Previously published data, in combination with our results strongly suggest that intradermal vaccinations generate long-lasting mucosal T-cell memory and neutralizing sIgA responses. In addition, due to the HPXV’s capacity to accommodate multiple antigens at various loci, HPXV-based vector vaccines may be able to elicit a more comprehensive immunity to targeted pathogens with a single vaccination. We believe the TNX-1800 construct can be modified to accommodate multiple SARS-CoV-2 genes such as Spike, Nucleocapsid protein (N), and the non-structural proteins (NSP) associated with the RNA-dependent RNA Polymerase. These strategies warrant further investigation.

Given that the TNX-1800 vaccine platform can induce a robust immune response with a single vaccination at small dose volumes, the vaccine is a viable candidate to be administered via microneedles. Further, the TNX-1800 vaccine platform is designed to enable single-dose, vial-sparing vaccines that can be manufactured using conventional cell culture systems, with the potential for mass-scale production, and packaging in multi-dose vials, that are not expected to require an ultra-cold-chain.

In summary, TNX-1800, is a live-replicating viral vaccine platform that provides induction of strong immunogenicity and efficacy after a single vaccination at low doses with good tolerability. The results are consistent with the idea that just a few rounds of replication generated efficacious immune responses against SARS-CoV-2 infection systemically and in the respiratory system. Collectively, these findings highlight the promising role of the TNX-1800 platform vaccine in advancing global efforts to halt the expansion of SARS-CoV-2 and other infectious diseases.

## Author Contributions

Conceptualization, A.M., B.D., S.J.G., H.S., S.L. and; methodology, J.P., F.K., R.N., S.J.G.; formal analysis, M.A., A.M., F.N., S.B., S.L.; investigation, M.A., A.M., F.N, S.B., S.L.; resources, A.M., F.N., S.B., S.L.; data curation, M.A., A.M., S.J.G., F.N., S.B.; writing—original draft preparation, M.A., S.B.; writing— review and editing, M.A., A.M., D.M., J.P., F.K., R.N., B.D., S.J.G., F.N., S.B., S.L.; visualization, M.A., A.M., F.N., S.B.; supervision, A.M., F.N., S.B., S.L.; project administration, A.M., S.B., S.L.; funding acquisition, S.L. All authors have read and agreed to the published version of the manuscript.

## Funding

This work was performed under a research agreement with Tonix Pharmaceuticals. The funder played a role in the formulation of the project, the decision to publish, and the preparation of the manuscript.

### Institutional Review Board Statement

This study was designed to use the fewest number of animals possible, consistent with the objective of the study, the scientific needs, contemporary scientific standards, and in consideration of applicable regulatory requirements. The study design was reviewed by the Institutional Animal Care and Use Committee (IACUC) at Southern Research and the study was in full compliance with applicable federal and Southern Research requirements. The Institutional Biosafety Committee (IBC) has approved this project. The facility where this research was conducted is accredited by the Association for Assessment and Accreditation of Laboratory Animal Care (AAALAC International) and adheres to principles stated in the Guide for the Care and Use of Laboratory Animals, National Research Council, 2011[11].

### Informed Consent Statement

Not applicable

### Data Availability Statement

Not applicable

## Acknowledgments

The authors wish to acknowledge Southern Research, BioQuol and members of Tonix Pharmaceuticals for providing insightful comments during the conduct of these studies.

